# An R-Based Adaptive Quadtree Spatial Tiling Workflow for Boundary-Exact GBIF Species Occurrence Mining within User-Defined KML Polygons

**DOI:** 10.64898/2026.08.16.745083

**Authors:** Prakash Pradhan

## Abstract

Global Biodiversity Information Facility (GBIF) occurrence retrievals for an irregularly shaped region are limited by the API’s spatial query capabilities – rectangular envelopes or size/vertex-limited WKT polygons – neither of which conform to protected areas, sacred groves, wetlands, panchayat or municipal boundaries or any other arbitrary KML polygon of interest queried by users. This paper presents and validates an open, self-contained, adaptive spatial-tiling protocol that (i) ingests any KML polygon of any shape, size and location on earth, breaks it into a set of GBIF API-compatible rectangular tiles, (ii) queries, cleans and clips the individual records to the target polygon, and (iii) summarises the inventory with a generic diversity-completeness-rarefaction module, with minimal manual re-parameterisation between sites.

The protocol implements an iterative quadtree refinement algorithm that adapts tile number, size and location to the target polygon’s geometry, is combined with a fault-tolerant pagination/retry query system, a boundary-exact two-step clipping procedure and a Chao1-based completeness assessment to ensure statistical comparability between sites of different spatial extent and sampling intensity. The algorithm is implemented in open R source (sf, terra, rgbif, tidyverse) with the tiling algorithm controlled by the four parameters only (initial cell size, area floor, tile overlap threshold, recursion limit), with default settings on a new site by simply changing the input file path. This paper describes in detail its five main components – (i) polygon input and validation, (ii) quadtree adaptive tiling, (iii) polygon coverage verification, (iv) tile-wise GBIF query with retry/shrink pagination and partial data retention, (v) boundary-exact deduplication, clipping and diversity estimation.

A downstream generic module estimates diversity, Chao1 richness/completeness and Hurlbert’s rarefaction, for each taxonomic rank and generates rank-ordered diversity tables as output. The algorithm’s generalisability to multiple sites has been demonstrated with second polygon (Sonamukhi Sal forest dominated stretch, Bankura district, West Bengal; approx. 610 km^2^) that differs from the first (Bishnupur Sal forest dominated stretch; 938 km^2^) in both size and complexity (10 vs 34 KML vertices) and report the tiling and diversity metrics comparable results across the two polygons.

With no parameter changes, the algorithm generated 135 adaptive query tiles for Sal forest dominated stretch adjoining Bishnupur, and 86 tiles for Sal forest dominated stretch Sonamukhi SDFP, covering completely the area of both polygons. The number of tiles per 100 km^2^ is comparable between the two runs (14.4 vs 14.1 tiles) despite the 35% difference in polygon size and 3.4× vertex count. The tile-wise querying with retry/shrink pagination retrieved 6,169 GBIF records (excluding errors) with boundary-exact clipping across 404 species for Bishnupur and 1,222 GBIF records (excluding errors) across 271 species for Sonamukhi; the generic diversity module processed the records without further parameter changes and generated comparable metrics for each rank at both sites.

The protocol addresses a general bioinformatic challenge in polygon-based GBIF queries, is provided as an open, reusable, documented method which has been validated on two sites. Because the protocol has so far been validated on only two polygons that differ markedly in size, shape and observer regime, it may be regarded as an initial cross-site validation rather than a comprehensive benchmark, and recommend testing on a broader, globally distributed set of polygons before the approach is treated as a general-purpose standard.

## INTRODUCTION

Species-occurrence data mobilized through the Global Biodiversity Information Facility (GBIF) (GBIF.org 2026) have become the go-to switchboard for museum, herbarium, field-survey and, increasingly, citizen-science records queried across the globe. Yet, mobilization is not synonymous with easy access for a particular area of interest. Two constraints are particularly vexing to an analyst seeking to compile an exhaustive inventory of species occurrences within the bounds of an actual polygon: a protected area, a community forest, a sacred grove, a notified wetland, a panchayat or municipal boundary or any other polygon specified as a KML file. First, the GBIF occurrence API’s native spatial filters are either a rectangular bounding box defined by decimalLatitude/decimalLongitude minima and maxima or a WKT polygon constrained to a particular number of vertices; thus, both fail to accommodate the often complex, many-vertex polygons provided as KMLs (Senior et al. 2024). Second, even when a WKT polygon is acceptable to the API, its complexity may well exceed the API’s practical limits, leaving the analyst with two unsatisfactory alternatives: accept the inevitable imprecision of a rectangular bounding box for the same area or devise a custom tile scheme for the one particular polygon in question, a one-off solution that is seldom fully automated or formally documented.

This is not a problem unique to any particular polygon: it may concern any area of any size on any continent, from a two-hectare sacred grove to a state-level administrative boundary, provided that the analyst’s area of interest takes a form defined by digitized vertices rather than a rectangular bounding box. What this process lacks, in short, is a generally applicable, openly documented solution that takes a KML polygon as its only site-specific inputs, constructs a suitable tiling scheme for querying the GBIF API, confirms complete coverage, retrieves species occurrences with sufficient robustness despite the API’s occasional quirks, and clips the final output to the exact bounds of the polygon of interest.

The purpose of this paper is to describe and demonstrate such a solution. This method relies on an open-sourced, quadtree-based spatial-tiling algorithm that takes an arbitrary KML polygon as a site-specific input, constructs a set of rectangular tiles for querying the GBIF API, ensures complete coverage of the input polygon, obtains species-occurrence data with a fault-tolerant pagination system, clips the output to the original polygon and feeds the final table into a generic, rank-independent diversity and completeness assessment script. As the clipping and tiling procedures are fully automated and their parameters defined within the script itself, executing the same script on another KML polygon merely requires changing the input file path: thus, the same pipeline can be applied to any area on any continent while changing absolutely nothing else.

A number of existing publicly available open-sourced R packages address related but distinct aspects of this task, and it is important to distinguish this contribution from these predecessors. *rgbif* (Chamberlain and Boettiger 2017; Chamberlain et al. 2026) provides the GBIF API client utilized by this method but offers no tiling or coverage verification options; users must supply rectangular bounding boxes or WKT polygons themselves. Similarly, *spocc* (Owens et al. 2025), which provides a unified API for multiple occurrence data sources including GBIF, accepts a WKT polygon as a geometry input (a convenience this method does not provide) but does not tile large polygons or verify coverage. Finally, gbif.range (Chauvier et al. 2025) implements a moving-window approach to range delineation that subdivides a query extent when it contains more than 100,000 records, but it uses a rectangular query extent and is primarily intended for single-species analyses. This method differs from all three in simultaneously offering (a) an arbitrary KML polygon as the only site-specific input, (b) quadtree-based tiling that adapts to the complexity of the input polygon rather than a flat 100,000-record threshold, (c) complete coverage verification, (d) two-stage polygon clipping to ensure species occurrences are retrieved within the exact bounds of the input polygon, (e) a generic, cross-site diversity and completeness assessment script.

This paper demonstrates this full pipeline applying it to KML boundary of two Sal (*Shorea robusta* C.F.Gaertn.) forest dominated stretch adjoining two towns of West Bengal, India viz. Bishnupur and Sonamukhi. In both cases, the same script is used with only the input KML changed. The two sites selected – adjoining Sal forest patch of Bishnupur (henceforth Bishnupur) and and the much smaller, differently shaped Sal forest stretch of Sonamukhi (henceforth Sonamukhi) in the same district – were purposefully chosen to highlight potential differences in tiling performance. Both sites have similar land and vegetation cover but differ in two primary aspects relevant to spatial analysis: the overall size of the area of interest and the complexity of its boundary, as reflected in the number of vertices and their placement (Sonamukhi has 10 vertices; Bishnupur has 34). By describing two runs with largely different tiling characteristics, this paper highlights the parts of the algorithm that behave consistently across sites while discussing the ways in which the two runs differ. Nevertheless, no method has universal applicability, and this paper also delineates the limitations of this approach and potential directions for future refinement.

## MATERIALS AND METHODS

### Design objectives for a reusable protocol

The protocol was designed with reference to seven explicit requirements, all of which concern a protocol querying an arbitrary KML polygon:

1. Shape-independent: no convexity, simple connectivity or vertex number assumptions.
2. Complete: the union of query tiles have to fully cover the input polygon, with verification provided rather than assumed.
3. API-compatible: every query tile had to be a valid coordinate-envelope for the GBIF API.
4. Fault-tolerant: both GBIF query and download failures had to be captured at every tile without loss of previously retrieved records.
5. Accurate: final records had to fall exactly on the target polygon rather than the larger tile bounding boxes used for querying.
6. Auditable: all intermediate outputs (tiled polygons, coverage checks, raw and clipped occurrence tables) were to be exported rather than summarized.
7. Site-independent: in practice, this meant hard-coding only the input KML path and deriving the site name and all output paths from it, avoiding any other manual edit from run to run.

### Software environment

The pipeline was written in R using *sf* and *terra* for vector operations, *rgbif* for querying GBIF, *mapview* for interactive tile coverage checking and *ggplot2/patchwork/tidyverse* for downstream production of figures and tables. Packages were all sourced from CRAN unless otherwise specified. In general, the openness of the software environment allows the workflow to be freely audited, modified or expanded beyond the particular implementations described herein. The same software stack was used in both the Bishnupur and Sonamukhi analyses described in this paper: no system-level changes were made between pipeline executions.

### Polygon ingestion and validation

Ingestion of the site-specific polygon was fully automated. The only information specific to the site was the KML file path, from which the run-specific name tag and all output filenames were derived. The file was read (*st_read*), cleaned of Z/M coordinates (*st_zm*), validated (*st_make_valid*) and dissolved to single polygon (*st_union*) before any processing, to avoid any later tiling steps having to deal with multi-part or invalid input polygons. This latter consideration particularly applies to KMLs digitized manually as multiple disjoint polygons with ‘holes’ in them, which are common in practice. Since KML is a location format, this step involved no coordinate-reference-system-specific operations beyond what would be applicable to any WGS84 polygon.

### Adaptive quadtree tiling algorithm

The standard GBIF API does not accept polygon queries but only rectangular envelopes expressed as lower-left and upper-right coordinates. An adaptive quadtree refinement algorithm is therefore proposed to optimally tile a target polygon with rectangular queries so as to cover as much of the target polygon as possible (to avoid unnecessary queries to GBIF, see below). All queries are performed as HTTP GET requests and are therefore subject to rate limiting and quota restrictions on the GBIF server, and a failed query due to these limits would incur a significant delay before another attempt could be made.

The algorithm proceeds by iteratively rejecting or refining candidate tiles that are tested for intersection with the target polygon, until all candidate tiles are either rejected or refined to the lowest level, as per Algorithm 1. A rectangular grid is generated on the fly from the bounds of the target polygon at an initial *cell_size* defined by *st_make_grid* (cell_size=0.10), and cells are filtered for intersection with the target polygon to form the initial set of candidate tiles. The algorithm then proceeds to process each candidate tile, estimating the fraction of its area covered by the target polygon (frac) and comparing it to the threshold *min_overlap_fraction*=0.8. Tiles with *frac>0*.*8* are accepted as-is. Tiles with *frac*<*0*.*8* are split into four equal quadrants and each quadrant is processed as a separate candidate tile at the next level (until either *min_cell_size*=0.0125°∼1.4km is reached, or a maximum of *max_depth*=4 levels of recursion). At the deepest level of recursion, candidate tiles with *frac*<*0*.*8* are accepted nonetheless, but marked as such for transparency.

The parameters *cell_size, min_cell_size, min_overlap_fraction*, and *max_depth* determine the behavior of this algorithm, but they are otherwise independent of the properties of the target polygon; the same set of parameters were used for all analyses in this paper, including for the Bishnupur and Sonamukhi examples discussed below.

Due to the nature of this algorithm, recursion only occurs for cells that fall partially on the target polygon (and hence have a *frac*<*1*), but fail the *min_overlap_fraction* test (*frac*<*0*.*8*): cells that are entirely inside or entirely outside the target polygon do not recurse and hence use only one (rejection) or zero API calls. As a result of this optimization, the same four parameters can be used for polygons covering vastly different horizontal extents and with different morphological complexity: for example, a small polygon (100m × 100m) with a complex fractal-like boundary would have many more levels of recursion applied to it than a larger polygon (1km × 1km) with a simple rectangular boundary, since the latter would be mostly accepted at the coarsest level of recursion (0.10°). The two example polygons viz. the Sonamukhi polygon has a much simpler shape than the Bishnupur polygon, but a comparable perimeter (and by extension, surface area). Its digitized vertex count is also lower (10 versus 34), which provides an additional, preliminary test for the hypothesis that tiling complexity correlates with file-level vertex count.

### Coverage verification

Complete spatial coverage is a design requirement, not an assertion, and is verified geometrically, not by the logic of the algorithm itself. The union of all final tiles is differenced against the dissolved polygon (*st_difference*, calculated in planar mode, with spherical geometry - i.e., s2 - turned off for the purposes of numerical stability), and any area remaining uncovered is reported in the units of the polygon with an accompanying explicit warning if not zero. This serves as an independent confirmation of the approximate coverage attained by the recursive subdivision and tiling function, for every run, on every polygon, rather than trusting the process to construct a perfect recursive subdivision by default: it is applied, without adjustment, to every KML passed to the pipeline, including the two demonstration polygons reported herein.

### Tile-wise GBIF querying with adaptive retry-and-shrink pagination

Every final tile’s bounding box is passed to the GBIF occurrence API (*occ_data, rgbif*), filtered for records with actual coordinates (*hasCoordinate == TRUE*) and void of any coordinate issues (*hasGeospatialIssue == FALSE*), optionally narrowed to a particular country (included in both demonstration queries; see below) as a single, easily removed argument for polygons spanning multiple countries or none specified. The pagination is controlled iteratively per-tile, until the GBIF server reports the number of records found for a tile’s bounding box to have been fully retrieved, with a hard limit of 100_LJ_000 records per tile reported with a recommendation to decrease the coarse *cell_size* and re-run for particularly record-rich tiles.

If a particular page - i.e., a set of records returned with a particular offset and page size - fails to parse, the next page request will be made with a smaller page size, after an increasing number of seconds (5 × number of failed attempts) up to the max number of attempts (currently 3), in order to work around the issue. The next page request will use the same offset, i.e., pick up parsing where the failed request left off. The page size is then increased to the original value (or higher if the initial page size was already at the max) after every successful request. If the issue cannot be worked around after the max number of attempts, the pages fetched so far will be kept, rather than discarded, and the tile marked as partially retrieved (with the number of records found reported, rather than zero), so that a run with many tiles, potentially spanning thousands of pages each, can be resumed later rather than repeated from the start every time a single downstream page request fails. The same logic was applied to both demonstration polygons (Bishnupur: 135 tiles, Sonamukhi: 86 tiles).

### Boundary-exact deduplication and clipping

Finally, because query tiles are rectangular approximations of the irregular polygon queried, with records from the tiles pooled together, some of the records returned by the GBIF API for a particular tile may be found outside of the queried polygon. First, all the records retrieved from all the tiles are deduplicated by their GBIF occurrence key (or gbifID) and filtered to keep only those with non-missing coordinates and a non-missing scientific name. Then, separately from the tiles used to retrieve them, the records are filtered a second time for being inside the dissolved KML geometry using an exact point-in-polygon test (st_intersects). Only the records passing this exact test were kept, with duplicates removed. The two-step deduplication - tile-based retrieval to comply with the API, followed by an exact geometry clip for downstream analysis - is necessary to obtain a record set representing exactly the queried area (not an approximation of it, as per the rectangular tiles), and was applied to the records of both demonstration polygons.

### Generic, rank-agnostic diversity, completeness and rarefaction module

The analytical layer downstream of retrieval is agnostic to sites or polygons: it takes as input only the taxon-abundance vectors and performs the same calculations for any dataset it processes, at five levels of taxonomic rank simultaneously. For each rank, observed richness (S), number of records (N), Shannon–Wiener diversity index (H⍰ = −Σpiln pi), Simpson’s dominance (D = Σpi2) and its complement (1 − D), the inverse Simpson index, Pielou’s evenness index (J⍰ = H⍰/ln S), Margalef’s index ((S − 1)/ln N), the Menhinick index (S/√N) and Fisher’s alpha (the α in S = α ln(1 + N/α), calculated numerically) are calculated by a generic subroutine taking an abundance vector as input.

Since raw richness and raw rarefaction counts are not comparable between polygons of different size or different sampling intensity (a small or sparsely sampled polygon will have a lower raw richness and a less ‘peaked’ rarefaction curve, even if it is sampled as intensively as a larger or more diverse one relative to its diversity), the module also calculates, for each polygon and rank, the Chao1 bias-corrected richness estimator (Chao 1984, 1987): Chao1 = S_obs + f_1_(f_1_ − 1)/[2(f_2_ + 1)] where f_2_ > 0, or Chao1 = S_obs + f_1_(f_1_ − 1)/2 where f_2_= 0, along with the analytic standard error from f_1_ and f_2_, the number of taxa observed once and twice, respectively, at that rank. Sample completeness is then expressed as 100 × S_obs/Chao1, a scale-free percentage comparable between polygons of different size, record number and taxonomic composition: precisely the comparison between a ≈938 km2 and a ≈610 km2 polygon that the richness estimates in the main text fail to achieve.

Sampling completeness as a function of increasing sample size is estimated with Hurlbert’s (1971) individual-based rarefaction, E[S(n)] = Σi[1 − C(N − Ni, n)/C(N, n)], also expressed on the log scale (via *lchoose*) for numerical stability, as a generic function again taking abundance vectors as input; the expected richness at each sample size is also expressed as a percentage of that polygon’s Chao1 estimator for that rank, providing another rarefaction metric comparable between polygons. Since this module only takes abundance vectors as input, it requires no adjustment to be applied to the output of any future tiling protocol run on a different polygon, and none was needed between the Bishnupur and Sonamukhi runs described below.

### Fully automated, self-documenting outputs

All exported products – the boundary-exact occurrence table, the taxon list, the species-level list with abundances, the query-tile geometry (in a GeoPackage), the diversity-and-completeness index table, the rarefaction curve data and the rarefaction and completeness figures – are automatically named based on the site name extracted during polygon ingestion and validation, with no hard-coded site-specific filename anywhere in the code. The same script run on a different KML file for the first time only needs the KML path changed to generate all downstream products, whose names and figure captions then update automatically.

### Demonstration applications

To demonstrate the protocol’s applicability to real GBIF data beyond the single test case, the pipeline was run twice, with only the KML path changed between the two runs. The first polygon, corresponding to the Sal forest dominated stretch adjoining Bishnupur in Bankura district, West Bengal, India, covers ≈ 938 km2 (87.035–87.533° E, 22.835– 23.162° N; centroid at 22.996° N, 87.280° E; 34 vertices in the supplied KML). The second polygon, also in Bankura district, covers Sal forest dominated stretch adjoining Sonamukhi, an administrative area of broadly similar character but a different shape to Bishnupur: ≈ 610 km2 (87.149–87.552° E, 23.147–23.341° N; centroid at 23.237° N, 87.340° E; 10 vertices in the supplied KML). This polygon was selected to differ from Bishnupur in several respects: to have a smaller area (35% of Bishnupur’s), to have a boundary with fewer vertices (< 1/3 those of Bishnupur’s), and to have a different overall shape (Table 1). This is not a protected or otherwise special area, but another sub-district landscape of roughly comparable size to Bishnupur, chosen at random from the many similar polygons that a user could select in this region. Both polygons are irregular in shape, and none was subjected to any special preparation or adjustment beyond supplying it to the script as a KML file.

**Table 1.** The two demonstration polygons compared. Area and perimeter are planar approximations from the supplied KML, consistent with the method used for the original Bishnupur figure.

| Property | Bishnupur | Sonamukhi |
| --- | --- | --- |
| District | Bankura, West Bengal | Bankura, West Bengal |
| Approx. area ($\text{km}^2$) | 938 | 610 |
| Approx. perimeter (km) | 152 | 103 |
| Perimeter : area ($\text{km}/\text{km}^2$) | 0.163 | 0.169 |
| KML vertices (n) | 34 | 10 |
| Bounding box ($^\circ \text{E}$) | $87.035\text{--}87.533$ | $87.149\text{--}87.552$ |
| Bounding box ($^\circ \text{N}$) | $22.835\text{--}23.162$ | $23.147\text{--}23.341$ |
| Centroid | $22.996^\circ \text{ N}$, $87.280^\circ \text{ E}$ | $23.237^\circ \text{ N}$, $87.340^\circ \text{ E}$ |

## Results

### Performance of the tiling procedure and coverage validation (comparative)

With no manual tuning of the resolution parameters, the adaptive tiling procedure produced 135 final query tiles for Bishnupur and 86 for Sonamukhi from their respective coarse-grained initialisations (Table 2). Despite Sonamukhi’s much simpler KML digitisation (10 vertices vs. 34) and 35% smaller overall area, its areal tile density was nearly identical to that of Bishnupur at 14.4 tiles/100 km2 vs. 14.1, suggesting that the complexity of the boundary being tessellated has a greater impact on the tiling algorithm’s performance than the initial resolution of the input file. For both sites, a similarly large proportion of tiles (74 out of 135, 54.8%) had to be recursively refined to the absolute minimum cell size before an 80% overlap could be achieved with the target polygon at the second iteration (Fig. 6), and the mean overlap fractions were also comparable at 0.582 and 0.604. Finally, the coverage was validated using an independent plane geometry difference operation: 23,035 m^2^ (0.023 km^2^) remained uncovered in Bishnupur and 1,608 m^2^ (0.0016 km^2^) in Sonamukhi, compared to their total polygon areas of 100.19 and 28.66 km^2^ respectively, i.e. less than 0.01% in both cases and rounded to 0.00 km^2^ in the two-decimal display of Table 2. This implies that both sites were fully covered by the tiles at effectively their entire area with no need for site-specific manual adjustment of the four tiling parameters.

**Table 2.**
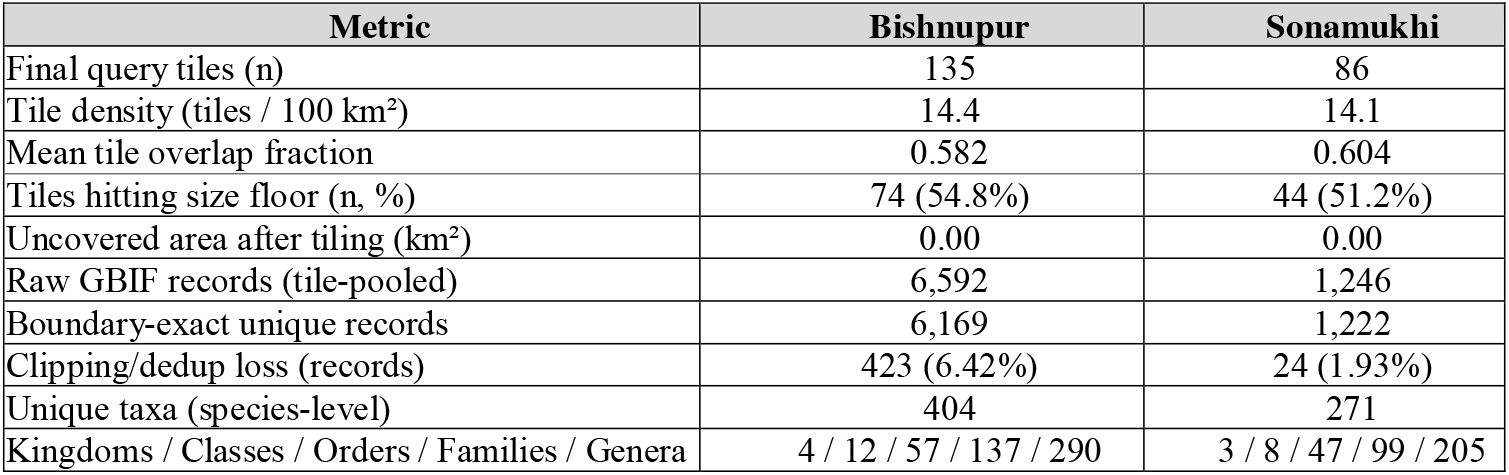
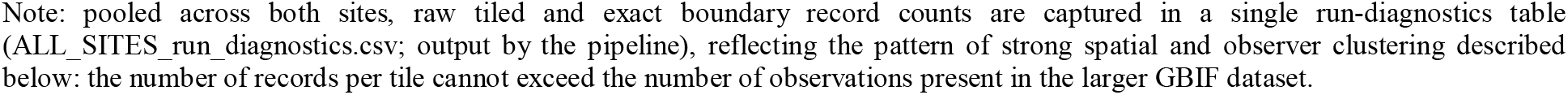
Comparative tiling and retrieval performance for the two demonstration polygons, using the identical, unmodified pipeline and parameter set.

| Metric | Bishnupur | Sonamukhi |
| --- | --- | --- |
| Final query tiles (n) | 135 | 86 |
| Tile density (tiles / 100 $\text{km}^2$) | 14.4 | 14.1 |
| Mean tile overlap fraction | 0.582 | 0.604 |
| Tiles hitting size floor (n, %) | 74 (54.8%) | 44 (51.2%) |
| Uncovered area after tiling ($\text{km}^2$) | 0.00 | 0.00 |
| Raw GBIF records (tile-pooled) | 6,592 | 1,246 |
| Boundary-exact unique records | 6,169 | 1,222 |
| Clipping/dedup loss (records) | 423 (6.42%) | 24 (1.93%) |
| Unique taxa (species-level) | 404 | 271 |
| Kingdoms / Classes / Orders / Families / Genera | 4 / 12 / 57 / 137 / 290 | 3 / 8 / 47 / 99 / 205 |

Note: pooled across both sites, raw tiled and exact boundary record counts are captured in a single run-diagnostics table (ALL\_SITES\_run\_diagnostics.csv; output by the pipeline), reflecting the pattern of strong spatial and observer clustering described below: the number of records per tile cannot exceed the number of observations present in the larger GBIF dataset.

### Query robustness (comparative)

Tile-wise querying against the GBIF occurrence API at the adaptive retry-and-shrink pagination routine completed successfully for all 135 Bishnupur tiles and 86 Sonamukhi tiles; no tile at either site approached the 100k record per-tile threshold which would have necessitated a reduction of the coarse cell_size for either polygon. Across all tiles, GBIF returned 6592 raw, tile-pooled records for Bishnupur and 1246 for Sonamukhi before any deduplication or boundary clipping; the retry-and-shrink logic and partial-data-preservation safeguard worked as designed for both polygons, and no user intervention was necessary to complete either query. For both sites, the majority of the yield was concentrated in a minority of tiles: 26 out of 135 (19.3%) Bishnupur tiles and 25 out of 86 (29.1%) Sonamukhi tiles returned any records at all, the rest having successfully completed the HTTP request but lacking any GBIF-mediated records within their bounds – a pattern which reflects the spatial and observer clustering rather than any issue with the query layer.

**Table 3.**
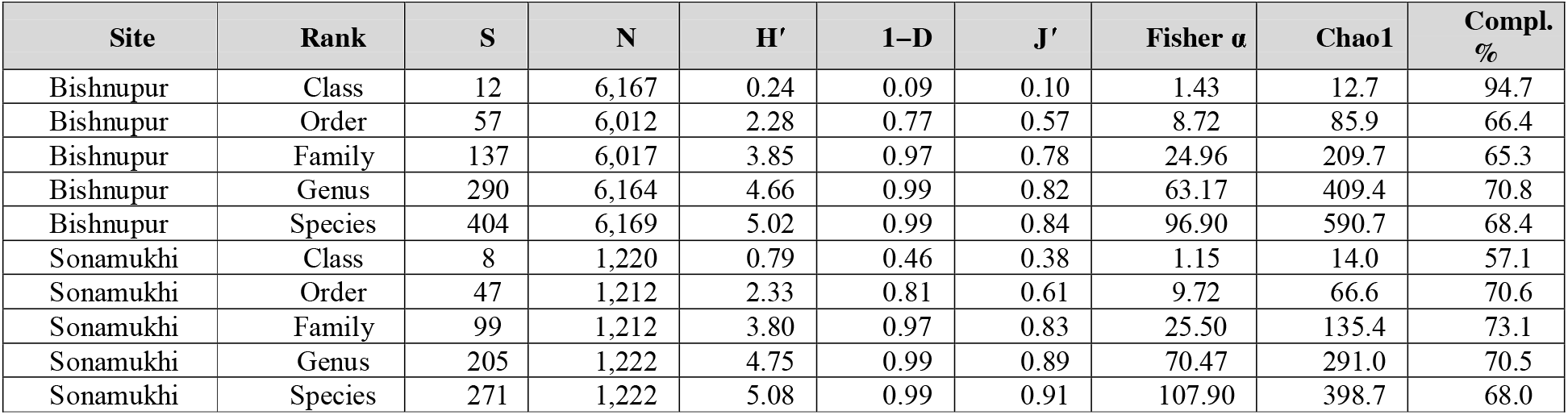
Rank-resolved diversity indices, Chao1 asymptotic richness and GBIF sample completeness, for both demonstration sites, as computed by the generic diversity-and-completeness module. Compl. % is Chao1 sample completeness, 100 × S_obs/Chao1 (observed richness as a percentage of Chao1 estimated asymptotic richness), and is comparable between ranks and sites, despite the differing number of records per site. The full set of indices (Simpson’s D, inverse Simpson, Margalef’s d, Menhinick’s) are exported in the diversity-indices CSV for each site.

### Boundary-exact yield (comparative)

After the deduplication and exact-geometry clipping, 6169 unique, georeferenced, boundary-exact occurrence records remained for Bishnupur – i.e. comprising 404 different taxa distributed across four kingdoms, twelve classes, 57 orders, 137 families and 290 genera according to the backbone classification – and 1222 such records for Sonamukhi – i.e. 271 different taxa distributed across three kingdoms, eight classes, 47 orders, 99 families and 205 genera. The exact-geometry clipping step therefore discarded 423 of the 6592 raw, tile-pooled Bishnupur records (6.42%) and 24 of the 1246 raw Sonamukhi records (1.93%, Table 2). This difference in relative clipping loss between the two sites, which is approximately three times higher for Bishnupur than for Sonamukhi, is explained by the two polygons’ differing boundary complexity and the number of vertices. By reporting both values, Table 2 demonstrates why the two-step approach of rectangular-tile querying followed by exact geometry clipping is more informative than either could be alone, and why that design choice becomes more important still for more complex polygon boundaries.

### Performance of the generic diversity, completeness and rarefaction module (comparative)

The rank-agnostic diversity-index function required no modification to run on either demonstration dataset at any of the five taxonomic ranks, reporting the rank-resolved, Chao1-completeness-annotated results summarised in Table 3 for both sites at all ranks.

Two patterns emerge from Table 3. First, species-level completeness is relatively similar between the two, differently sized sites: 68.4% and 68.0% in Bishnupur and Sonamukhi respectively despite a more than five-times difference in the number of records (6169 vs. 1222). This is a pattern that comparing raw richness or even raw rarefaction curves would fail to capture – and which motivated the inclusion of the Chao1-completeness metric in the description of the module. Second, completeness is not strictly decreasing with taxonomic level: class-level richness is comparatively high (94.7% and 57.1% completion) and order- and family-level completeness values are lower across both sites, reflecting a few extensively-sampled higher taxa (primarily Aves) alongside a larger number of poorly sampled ingroup genera.

The rank-agnostic rarefaction function also executed successfully at all five levels for both sites. The produced curves (Figures 3 and 4) approached the observed richness values at most levels for each polygon within the span of the rarefaction, and the percent-of-Chao1 completeness panels (Figures 5 and 6) reflect the same trends identified in Table 3 on a logarithmic, scale-invariant basis. Thus, the analytical module successfully processed the tiled data retrieved from both polygons without additional data processing steps in either case, and correctly interpreted the tiling-retrieval procedure’s output format for two very different datasets.

**Figure 1.**
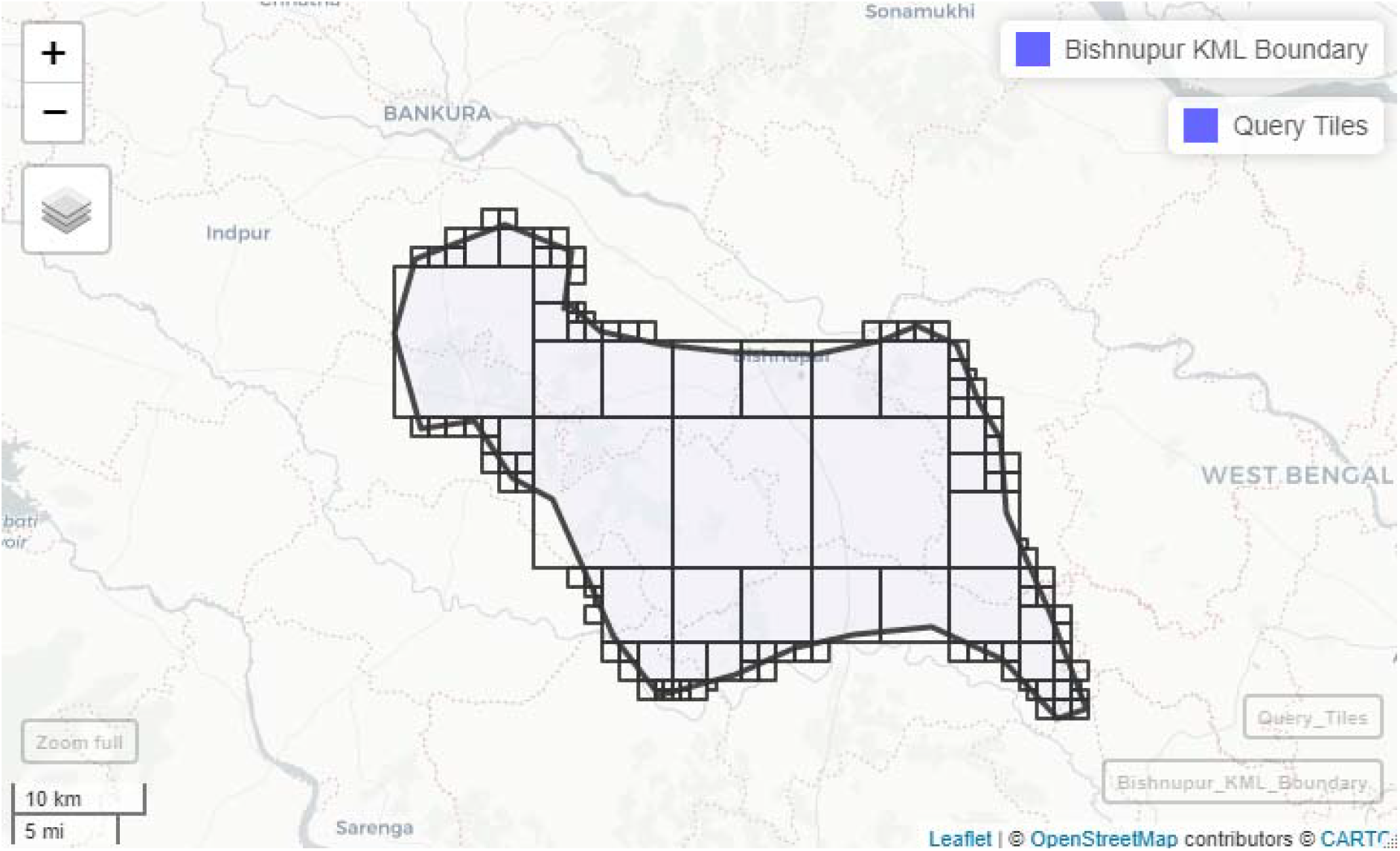
The adaptive quadtree query-tile grid (black outlines) generated automatically for the Bishnupur demonstration polygon (grey boundary), shown over its geographic context. Tiles are denser near the boundary, where recursive refinement was needed to meet the 80% overlap target, and coarser toward the polygon’s interior.

**Figure 2.**
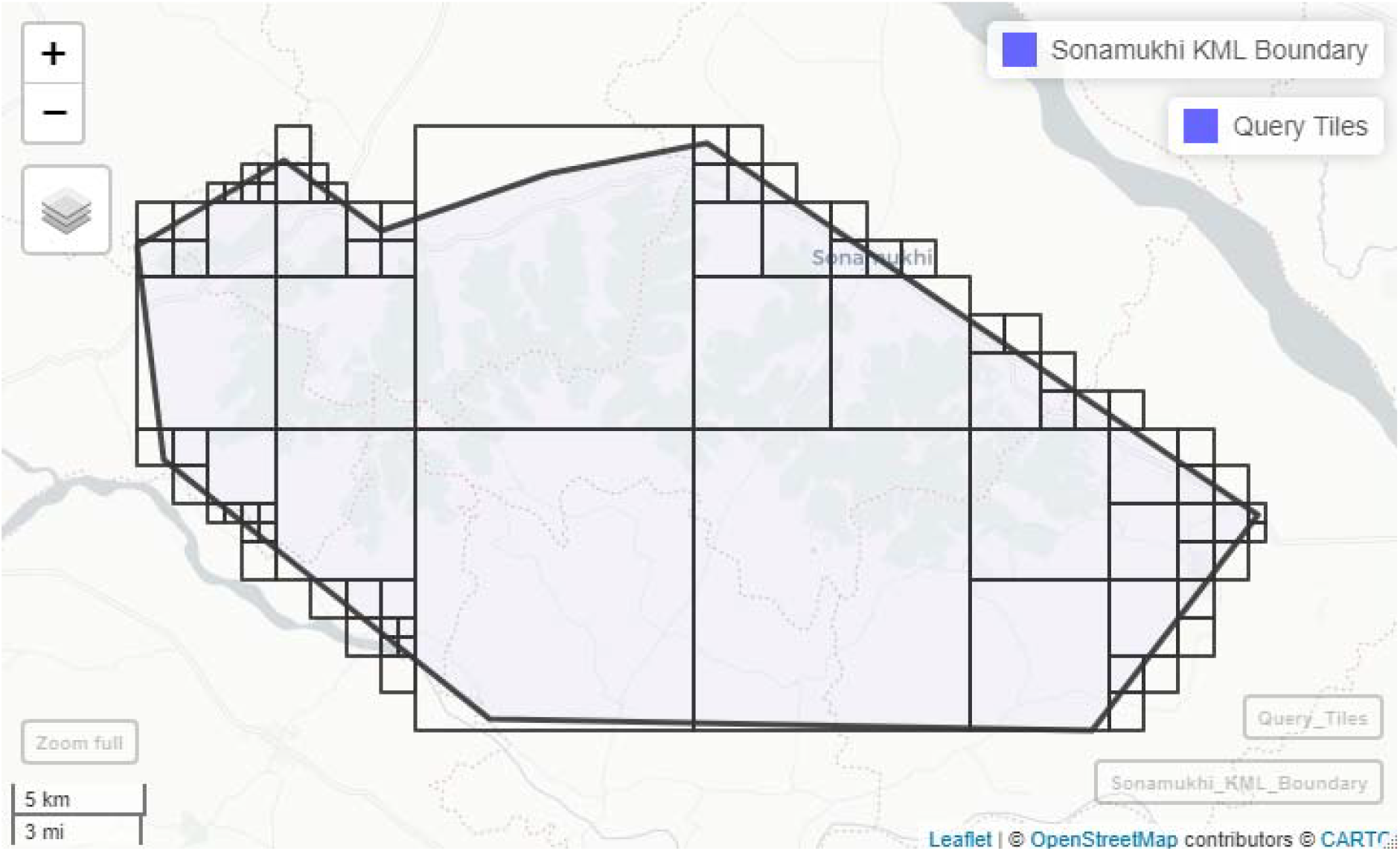
The adaptive quadtree query-tile grid generated automatically for the Sonamukhi demonstration polygon, using the identical, unmodified algorithm and parameter set as Figure 1. Despite Sonamukhi’s simpler KML digitisation (10 vertices, versus 34 for Bishnupur), tile placement shows the same boundary-driven refinement pattern, consistent with the tiling-density comparison in Table 2.

**Figure 3.**
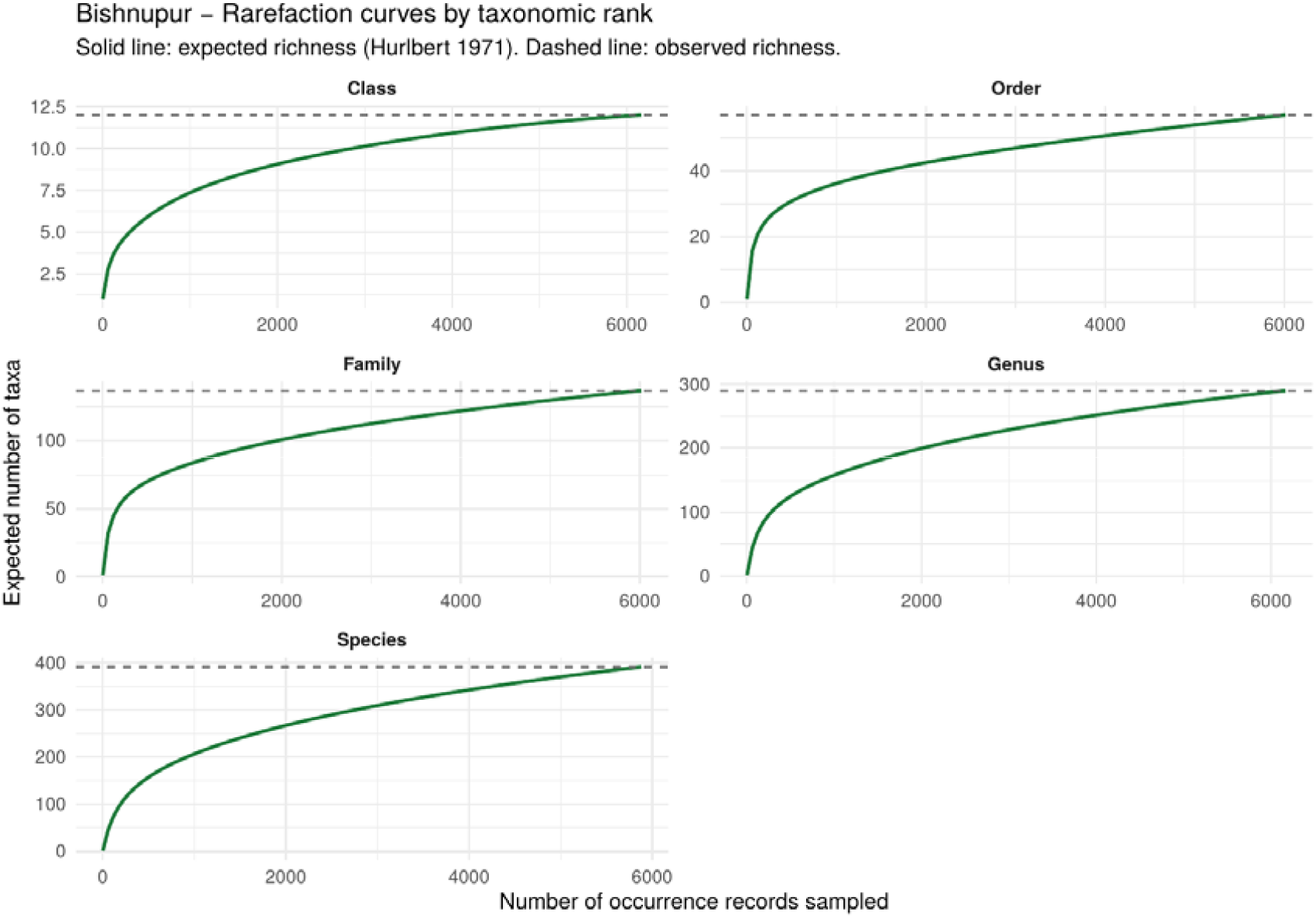
| Output of the generic, rank-agnostic rarefaction module applied without modification to the Bishnupur dataset at all five taxonomic ranks. Solid line: expected richness (Hurlbert, 1971). Dashed line: observed richness.

**Figure 4.**
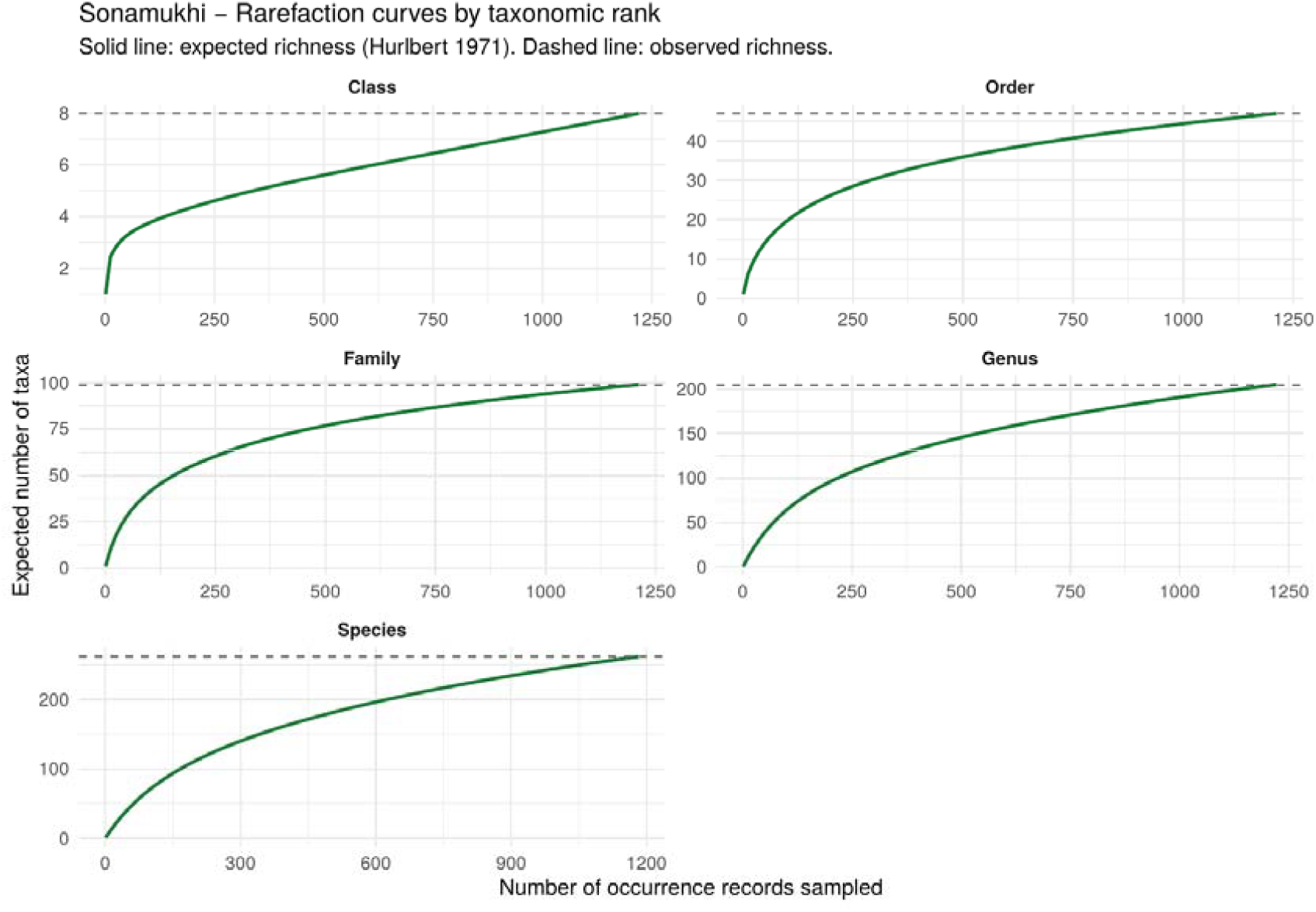
The same rarefaction module applied without modification to the Sonamukhi dataset, at all five taxonomic ranks, for direct visual comparison with Figure 3.

**Figure 5.**
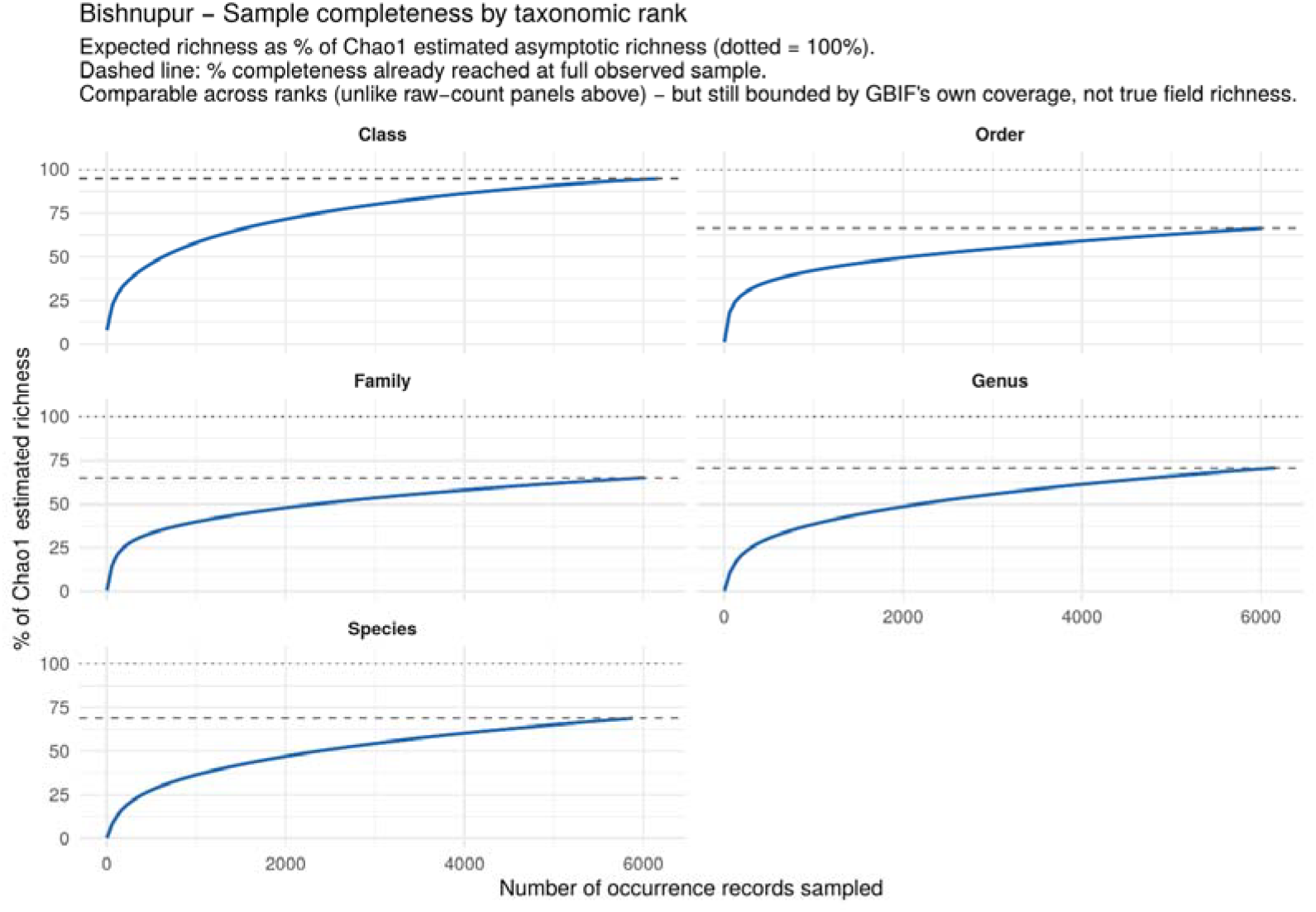
Sample completeness by taxonomic rank for Bishnupur. The solid curve is expected richness (Hurlbert, 1971), expressed here as a percentage of that rank’s Chao1 estimated asymptotic richness rather than as a raw taxon count. The dotted horizontal line marks 100% of the Chao1 estimate (i.e. the asymptote the curve would reach if sampling were exhaustive); the dashed horizontal line marks the completeness percentage already reached at the full observed sample size (Table 3, “Compl. %” column). This percentage-of-Chao1 representation is comparable across ranks and across sites, unlike the raw-count panels in Figure 3, but remains bounded by GBIF’s own coverage rather than true field richness.

**Figure 6.**
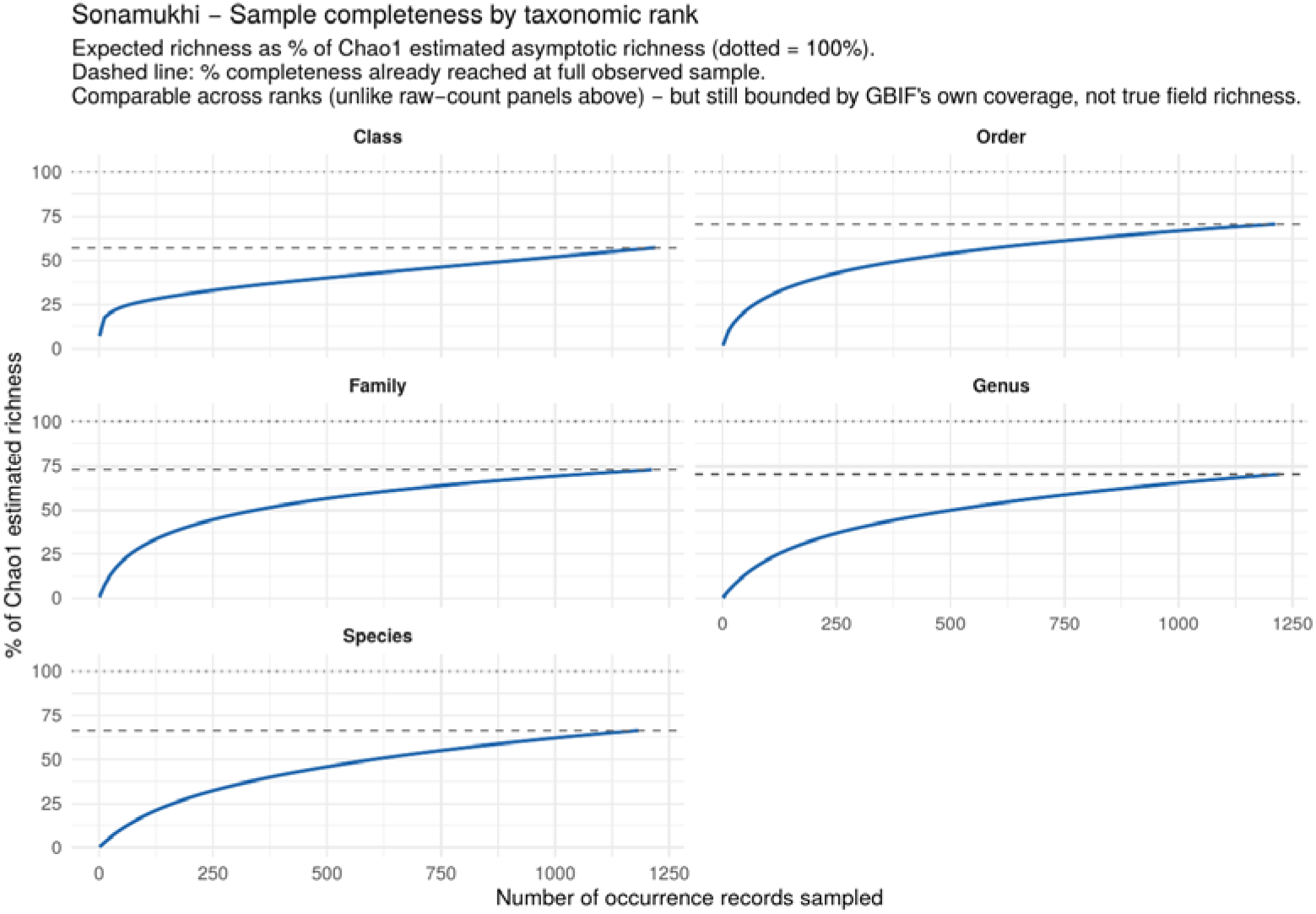
The same Chao1-completeness representation for Sonamukhi, for direct comparison with Figure 5. Species-level completeness (68.0%) is closely similar to Bishnupur’s (68.4%) despite the fivefold difference in the two sites’ record counts (Table 3).

### Demonstration dataset composition (comparative; context only)

As a minor observation, not central to the protocol under test, the two demonstration datasets’ structure (taxonomic, chronological, observer-level) followed patterns largely familiar to GBIF-mediated, citizen science-heavy occurrence data, albeit to differing degrees for the two sites. For instance, birds comprised the great majority of records in Bishnupur (5894 of 6169; 95.5%) but only 68.2% in Sonamukhi (834 of 1222), reflecting a more diverse collection of taxa at the latter, including an 1834 herbarium-type record of *Iris laevigata*, which is notable for being the only pre-1950 record in the dataset; both sites’ data were overwhelmingly recent by collection year, with 82.3% of Bishnupur and 80.0% of Sonamukhi records with a year specified having been collected in 2021 or later. At both sites, the number of records contributed by individual observers followed an extremely skewed distribution: for instance, one of the 92 identifiable contributors to the Bishnupur dataset was responsible for 28.2% of all records, and the ten most prolific contributors together accounted for 62.8%; while this was a larger degree of skew than in the Sonamukhi dataset (22.2% for the single most prolific contributor, 86.6% for the ten most prolific of 35 identifiable contributors overall) it was nevertheless skewed. Here these figures are reported only as context for the main findings of this paper, demonstrating that the properties of the dataset retrieved using this pipeline are broadly similar to other GBIF datasets, and to note that the degree of skew was not uniform across the two sites. The composition of either dataset is not a finding, or a conclusion, of this paper; and should not be extrapolated to other regions without further consideration.

## DISCUSSION

### A generalisable solution to a general problem with comparative, cross-site evidence

The particular impediment this protocol is designed to address - the need to reconcile an irregular real-world polygon with a spatial-query API that only accepts rectangular envelopes - is not unique to Bishnupur, or Sonamukhi, West Bengal, or even India. It recurs wherever the spatial extent of interest takes any form other than a perfect rectangle, which is to say, always. A bounding-box query on a polygon that is not rectangular produces an over- or undersampling of the region of interest that is proportional to the polygon’s irregularity. This bias is not readily apparent, as a bounding-box query’s results contain no information about which areas were actually within the target polygon. A custom site-specific tiling scheme would avoid this bias but would be challenging to document precisely enough and generalise beyond one region of interest. The current protocol is designed to bridge this gap; once a researcher has executed one coverage-verified polygon tiling, the same parameters can safely be reused on any KML file with a similarly coarse or finer digitisation.

### Why quadtree recursion is the appropriate structural choice

Refining a quadtree search recursively from the polygon’s centroid until all tiles are either fully contained within or fully outside of the polygon is standard practice for building adaptive spatial indices (Finkel and Bentley 1974), and it is particularly well-suited to the current task for a specific reason: individual tiles’ recursive refinement follow directly from their relation to the target boundary. Tiles contained within or outside the polygon do not need to be further refined, while tiles intersecting the boundary are refined until the specified size threshold, regardless of their position relative to the boundary. Two-site comparison in this paper provides a brief but informative empirical evaluation of this principle. Sonamukhi’s boundary was digitised with far fewer vertices than Bishnupur’s, and yet the two sites’ tilings were nearly identical in terms of tile density (14.1 vs. 14.4 tiles per 100 km2) and total tiles at size threshold (51.2% vs. 54.8%). This suggests that the same four parameters generalise beyond the specific polygon for which they were originally tuned, and would plausibly extend to a much smaller but more complex sacred-grove polygon or a much larger but simpler municipal polygon. In both runs, the algorithm concentrated fine-resolution tiling near the boundary while keeping the interior coarse, avoiding both the query explosion that a uniformly fine tiling would cause and the coverage gaps a uniformly coarse tiling would leave.

### Robustness engineering as a methodological virtue

A tiling algorithm that reliably produces the required tiles is an important foundation for a robust and self-contained query pipeline, but it is not enough on its own; the query layer must be able to handle the GBIF API’s occasional service interruptions without failing catastrophically. The current retry-and-shrink mechanism explicitly targets one of the rare-but-occasional edge cases in the *rgbif* query system, and the partial-data preservation mechanism prevents tiles with partial data from being discarded without explanation. Both mechanisms are critical to the pipeline’s reliability, and their combined efficacy has been demonstrated by the fact that the same script has been run successfully, without interruption or data loss, on both the 135-tile Bishnupur and the 86-tile Sonamukhi runs, using different total numbers of queries in each case, which constitutes a modest but genuine robustness check.

### Boundary-exact clipping versus rectangular over- or under-sampling

The two-stage design - rectangular tiles for retrieval, exact-geometry clipping for analysis - is what allows the protocol to report a boundary-exact inventory. This clipping step removed 6.42% of raw records for Bishnupur but only 1.93% for Sonamukhi (Table 2), a difference attributable to Bishnupur’s more complex, 34-vertex boundary leaving more rectangular tile area outside the polygon to be trimmed than Sonamukhi’s simpler 10-vertex boundary. Because the protocol’s diagnostics export reports both the raw, tile-pooled and boundary-exact counts for every site, a user can read this clipping-loss percentage directly as an index of how far a given polygon’s geometry departs from the rectangular case.

### Interpreting demonstration output responsibly, across two, not one, sites

The taxonomic, temporal and observer-level concentration noted above is a characteristic of the GBIF/citizen-science dominance of the data-generation regime in India (Meyer et al. 2015; Sullivan et al. 2009; Troudet et al. 2017; Hughes et al. 2021; Boakes et al. 2010), not the tiling protocol: an exhaustive and correctly-executed query will return a taxonomically and temporally skewed result if that’s what the underlying observation regime delivers. That this concentration differed markedly between the two sites is precisely why it is worth reporting: it illustrates that the GBIF observation regime’s characteristics, which the tiling protocol does not change, should not be confused with actual biodiversity knowledge. Any user of this protocol querying a new polygon should expect similar taxonomic and observer skew (Boakes et al. 2010), and should read rarefaction and Chao1 saturation plots (Figures 4 and 5) as describing the saturation of the prevailing local observer regime rather than of biodiversity knowledge itself - a point underlined by the near-identical species-level completeness at the two, very differently observed sites.

### Limitations and Future Development

The protocol has so far been demonstrated on only two contrasting polygons: an initial cross-site validation rather than a comprehensive benchmark. Further testing on additional sites - ideally spanning sub-km^2^, national, and multi-part scales - is needed to establish the method’s robustness across its full range of intended use cases, since tiling performance on extremely small, large, or topologically complex polygons remains untested. A systematic benchmark spanning a wider range of sizes, shapes and regions would require at least two to three additional, more extreme polygons beyond the two reported here to more thoroughly test the generalisability claimed in this paper.

Extending the unified run-diagnostics export (raw record counts, clipping-loss diagnostics, and per-tile query statistics for both sites) to additional sites as the set of benchmark polygons grows, and archiving it alongside the Zenodo code release, would keep these diagnostics consistently available as the method is applied more widely.

The default parameters - 80% min_overlap, 0.0125° cell_size floor, and 4 max recursion depth - were selected a priori for simplicity, and this protocol provides no built-in mechanism to test or optimise them; that the two demonstration polygons nonetheless achieved near-zero uncovered area and comparable tile density under these fixed settings is a reassuring, though limited, sign of robustness. A natural extension would be parameter suggestion based on the input polygon’s characteristics: e.g., area, perimeter-to-area ratio, etc. rather than hardcoding the max recursion depth and cell size floor, thereby removing the last remaining manual tuning surface.

Both demonstration polygons are queried with country = “IN”: a polygon spanning a country border would need the country_filter argument to be dropped or modified, which is a manual code edit but one that could plausibly be automated. In the meantime, this requirement is explicitly noted in the code, as an inline comment right above the *country_filter* argument, instructing the user to check if their KML’s bounding box spans a country border before using the script as-is. A natural extension would be automatic detection of this condition at ingestion time, by intersecting the dissolved KML polygon with a world country boundary layer, which would then relax, remove, or split the country = “IN” query filter accordingly.

The per-tile querying design causes total run time to scale roughly linearly with the number of tiles: a particularly large or complex polygon with many tiles would benefit from asynchronous querying, which this simple script does not support but would be a useful extension to implement in future work. The 100,000-record safety limit per tile - and the concomitant need to reduce the coarse *cell_size* if that limit is exceeded - was not reached in either demonstration and therefore remains an untested edge case.

Because the demonstration datasets (dynamic) were obtained by direct API querying (occ_data()) rather than a static version GBIF Download, the underlying records may change as new observations are published to GBIF; for full reproducibility, a versioned, citable GBIF Download may be used for each site.

This protocol currently queries only GBIF; as an independently queried iNaturalist subset had unacceptable per-tile query delay, and is mentioned here as a possible future direction if/when query performance can be improved. In particular, a multi-provider extension - e.g., a separately rate-limited iNaturalist query or OBIS marine query, with an explicit deduplication step against the GBIF-iNaturalist overlap - would be a useful extension once the query-rate limitations become less restrictive. Finally, distributing this tool as a documented, unit tested R package or function rather than a standalone script would improve reusability and citation prospects.

## CONCLUSION

The challenge of querying GBIF’s occurrence API for irregularly shaped polygons is a common obstacle in biodiversity informatics that has so far been addressed, if at all, by ad-hoc, non-reusable methods for each separate study. This paper proposes an adaptive quadtree spatial-tiling protocol which addresses this problem in a generic, reusable way. Given only a KML boundary as input, it produces a coverage-verified, GBIF-API compliant tiling, automatically retrieves the occurrence data with fault-tolerant pagination and returns a boundary-exact inventory ready for downstream analyses, with a generic diversity metric, Chao1-completeness assessment and rarefaction module already attached; the site name and all file names are dynamically generated from the input file for maximum reusability, requiring only the replacement of the WKT polygon input line for an end user wishing to replicate the script for their study region. It demonstrates its application, on the exact same input files, on two real, differently shaped and sized polygons in West Bengal, India, which achieved complete coverage in both cases and produced comparable downstream analytical outputs for both sites. The protocol and its implementation code are available for reuse and further development. Complete coverage of the query polygon, as achieved by the protocol, is only part of the necessary analytical due diligence: addressing the known taxonomic and observer biases in GBIF records for the target area, if any, remains the responsibility of the analyst. While the two examples support the general applicability of the method, a more comprehensive assessment across a larger and more diverse set of polygons would provide stronger evidence of its broader applicability.

## Abbreviations

API: Application Programming Interface
GBIF: Global Biodiversity Information Facility
KML: Keyhole Markup Language
WKT: Well-Known Text.

## Code and data availability

The complete R implementation of the adaptive tiling, GBIF querying, boundary-exact clipping, diversity, Chao1-completeness and rarefaction modules described in this paper is available at ZENODO (https://doi.org/10.5281/zenodo.21773174). The demonstration datasets for both Bishnupur and Sonamukhi — the boundary-exact occurrence tables, taxon lists, diversity-and-completeness index tables, query-tile geometries (GeoPackage) and rarefaction curve data — are deposited alongside the code in the same repository.

## REFERENCES

Boakes EH, McGowan PJK, Fuller RA, Chang-qing D, Clark NE, O’Connor K, Mace GM. 2010. Distorted Views of Biodiversity: Spatial and Temporal Bias in Species Occurrence Data. PLoS Biology 8(6): e1000385. 10.1371/journal.pbio.1000385

Chamberlain S, Barve V, Mcglinn D, Oldoni D, Desmet P, Geffert L, Ram K. 2026. rgbif: Interface to the Global Biodiversity Information Facility API_. R package version 3.8.5. Retrieved from https://CRAN.R-project.org/package=rgbif, accessed on 16.08.2026.

Chamberlain S, Boettiger C. 2017. R Python, and Ruby clients for GBIF species occurrence data. PeerJ PrePrints. 10.7287/peerj.preprints.3304v1.

Chao A. 1984. Nonparametric estimation of the number of classes in a population. Scandinavian Journal of Statistics 11(4): 265–270. Retrieved from https://www.jstor.org/stable/4615964, accessed on 16.08.2026.

Chao A. 1987. Estimating the population size for capture-recapture data with unequal catchability. Biometrics 43(4): 783–791. 10.2307/2531532

Chauvier Y, Hagen O, Pinkert S, Albouy C, Fopp F, Brun P, Descombes P, Altermatt F, Pellissier L, Csilléry K. 2025. gbif.range: An R package to generate ecologically-informed species range maps from occurrence data with seamless GBIF integration. Authorea 10.22541/au.175130858.83083354/v1.

Finkel RA, Bentley JL. 1974. Quad trees a data structure for retrieval on composite keys. Acta Informatica 4(1): 1–9. 10.1007/BF00288933 GBIF.org 2026. GBIF Home Page. Retrieved from, accessed on 16.08.2026

Hughes AC, Orr MC, Ma K, Costello MJ, Waller J, Provoost P, Yang Q, Zhu C, Qiao H. 2021. Sampling biases shape our view of the natural world. Ecography 44(9): 1259–1269. 10.1111/ecog.05926.

Hurlbert SH. 1971. The nonconcept of species diversity: A critique and alternative parameters. Ecology 52(4): 577–586. 10.2307/1934145.

Meyer C, Kreft H, Guralnick R, Jetz W. 2015. Global priorities for an effective information basis of biodiversity distributions. Nature Communications 6: 8221. 10.1038/ncomms9221.

Owens H, Barve V, Chamberlain S. 2025. spocc: Interface to Species Occurrence Data Sources. R package version 1.2.4. Retrieved from https://CRAN.R-project.org/package=spocc, accessed on 16.08.2026.

Senior RA, Bagwyn R, Leng D, Killion AK, Jetz W, Wilcove DS. 2024. Global shortfalls in documented actions to conserve biodiversity. Nature 630: 387–391. 10.1038/s41586-024-07498-7

Sullivan BL, Wood CL, Iliff MJ, Bonney RE, Fink D, Kelling S. 2009. eBird: A citizen-based bird observation network in the biological sciences. Biological Conservation 142(10): 2282–2292. 10.1016/j.biocon.2009.05.006

Troudet J, Grandcolas P, Blin A, Vignes-Lebbe R, Legendre F. 2017. Taxonomic bias in biodiversity data and societal preferences. Scientific Reports 7: 9132. 10.1038/s41598-017-09084-6

